# Algorithms for Estimating Time-Locked Neural Response Components in Cortical Processing of Continuous Speech

**DOI:** 10.1101/2022.01.18.476815

**Authors:** Joshua P. Kulasingham, Jonathan Z. Simon

## Abstract

**Objective:** The Temporal Response Function (TRF) is a linear model of neural activity time-locked to continuous stimuli, including continuous speech. TRFs based on speech envelopes typically have distinct components that have provided remarkable insights into the cortical processing of speech. However, current methods may lead to less than reliable estimates of single-subject TRF components. Here, we compare two established methods, in TRF component estimation, and also propose novel algorithms that utilize prior knowledge of these components, bypassing the full TRF estimation.

**Methods:** We compared two established algorithms, ridge and boosting, and two novel algorithms based on Subspace Pursuit (SP) and Expectation Maximization (EM), which directly estimate TRF components given plausible assumptions regarding component characteristics. Single-channel, multi-channel, and source-localized TRFs were fit on simulations and real magnetoencephalographic data. Performance metrics included model fit and component estimation accuracy.

**Results:** Boosting and ridge have comparable performance in component estimation. The novel algorithms outperformed the others in simulations, but not on real data, possibly due to the plausible assumptions not actually being met. Ridge had slightly better model fits on real data compared to boosting, but also more spurious TRF activity.

**Conclusion:** Results indicate that both smooth (ridge) and sparse (boosting) algorithms perform comparably at TRF component estimation. The SP and EM algorithms may be accurate, but rely on assumptions of component characteristics.

**Significance:** This systematic comparison establishes the suitability of widely used and novel algorithms for estimating robust TRF components, which is essential for improved subject-specific investigations into the cortical processing of speech.

## I. Introduction

**T**HE human brain time-locks to features of continuous speech, extracting meaningful information relevant to comprehension. Magnetoencephalography (MEG) and electroencephalography (EEG) are suitable methods to measure these time-locked responses, due to their high temporal resolution. Traditional methods for analyzing auditory responses involve averaging over multiple trials of repeated stimuli to estimate Evoked Response Potentials (ERPs) [1], [2]. But exploring the complex mechanisms involved in speech processing requires non-repetitive, continuous speech stimuli of long duration, and averaging over trials is no longer feasible. One method of analyzing responses to continuous stimuli uses linear models called Temporal Response Functions (TRFs), that estimate the impulse response of the neural system to continuous stimuli [3], [4]. TRFs based on neural recordings using magnetoencephalography (MEG) have response components such as the M50 (~50 ms latency), M100 (~100-150 ms) and M200 (~200-250 ms) that are analogous to well-known auditory ERP components, the P1, N1, and P2 components of electroencephalography (EEG), and which have been utilized to investigate selective attention [3], [5]–[7], linguistic processing [8]–[10], and age-related differences in the auditory system [11]. However, though estimated TRFs display these canonical components at the group-average level, individual TRFs are much noisier and do not always have well-defined components. It is essential to detect robust response components on a per-subject level, both to identify task effects and for biomedical applications such as smart hearing aids. Hence, the suitability of various TRF methods for component estimation must be determined.

Variations of regularized regression and machine learning methods for estimating TRFs have been previously compared for decoding subject attention in a multi-talker scenario [6], [12], [13]. However, it is unclear how they compare to commonly used sparse TRF estimation techniques such as boosting [14], [15]. Furthermore, a focus on model fits for attention decoding may not be suitable for studies interested in accurate estimation of TRF components.

In this work we perform a systematic comparison of TRF algorithms in terms of estimating TRF components. Two widely used TRF estimation algorithms are ridge regression [13], [16] and boosting [3], [14], [15]. The former uses ℓ_2_ regularization which leads to smooth TRFs with broad components, while the latter greedily adds values to the TRF, thereby prioritizing sparsity in the TRF and leading to narrower, sharper components. However, it is not clear which of these methods is more accurate in estimating TRF component latencies and amplitudes.

Both ridge and boosting do not place restrictions on the number or latencies of specific TRF components. Since canonical auditory response components are often present in TRFs to the speech envelope, it is reasonable to incorporate this information during estimation. Several methods have been proposed for directly estimating latencies and amplitudes for M/EEG evoked responses (but not for TRFs). The earliest ERP latency estimation methods involved cross correlation with average response templates [17]. More recent algorithms have utilized techniques such as Independent Component Analysis [18], [19], wavelet decomposition [20], maximum likelihood estimation [21], [22], autoregressive models [23], Expectation Maximization (EM) [24], Matching Pursuit [25] and Bayesian methods [26], [27].

In this work, we propose novel TRF component estimation algorithms that utilize prior knowledge of the characteristics of neural responses (i.e., component latency ranges), and directly estimate component latencies, amplitudes and topographies. The first proposed algorithm estimates single-channel TRF component latencies and amplitudes using Subspace Pursuit (SP) [28]. The second algorithm extends this method for multi-channel TRFs using SP and Expectation Maximization (EM) [24], [29], and also directly estimates sensor topographies or cortical source distributions of TRF components. The SP algorithm is widely used for sparse signal recovery and is typically capable of recovering components in an efficient manner. The EM algorithm is a maximum likelihood method that is able to incorporate ‘hidden’ variables and is widely used in signal estimation [30]. Pursuit algorithms and EM have been used for single trial evoked response estimation [24], [25], and here, we employ natural extensions of these algorithms for TRF component estimation.

A simulation study, and an application of these algorithms to a real dataset, are reported and their performance is compared using single-channel, multi-channel, and source localized TRFs. Performance metrics include the correlation between the actual and the predicted signal, which is the conventional measure of model fit, and several other measures of component estimation accuracy. Throughout this work, “model fit” denotes the Pearson correlation between the actual and predicted signals. Other considerations such as spurious TRF activity and missing components are also examined. In summary, this work discusses the strengths and weaknesses of widely used algorithms and proposes novel methods for TRF component estimation that may provide robust and interpretable time-locked response components.

## II. Methods

### A. Established Algorithms for TRF estimation

The TRF estimation problem is given by the convolution

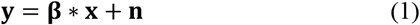

Where 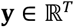 is the vector of the single-channel measured signal (e.g., at one sensor) for *T* time points, 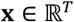 is the predictor variable (e.g., the speech envelope), 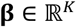 is the corresponding TRF over *K* time lags, and 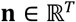 is the noise. This can be reformulated as a regression as follows

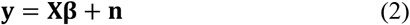

Where 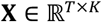 is the Toeplitz matrix formed by lagged predictor values. The well-known ridge regression algorithm has been widely used to solve this problem [16]. Another commonly used technique is the boosting algorithm, which is a sparse estimation technique belonging to the broad family of greedy additive estimators, and solves the TRF problem using coordinate descent [14], [15], [31]. In brief, this algorithm starts from an all-zero TRF and incrementally adds small, fixed values to the TRF to decrease the mean square error (MSE) at each iteration. The iterations are stopped when the MSE does not improve. A dictionary of basis elements (e.g., Hamming windows) is used for the incremental additions to the TRF. Both ridge and boosting can be used independently at each sensor to estimate TRFs for multi-channel data.

### B. Proposed SP algorithm for TRF estimation

The SP algorithm searches for TRF components within predefined latency windows and directly estimates them. This is unlike the ridge and boosting algorithms that do not place specific restrictions on the number or latencies of detected TRF components. Assuming there are *J* components (e.g., *J* = 3 for M50, M100, M200 components), the TRF model is now given by a modified version of (1).

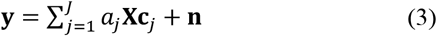

Where 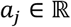 and 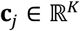 are the amplitude and waveform for the *j^th^* component. The component waveforms **c***_j_* are selected according to the component latency *τ_j_* from a basis dictionary (e.g., Hamming windows) that span the TRF lags (i.e., **c***_j_* is column number *τ_j_* of the basis dictionary matrix). The SP algorithm directly estimates the amplitudes *a_j_* and latencies *τ_j_*. The complete algorithm is given in Algorithm 1.

#### Algorithm 1: SP for TRF estimation

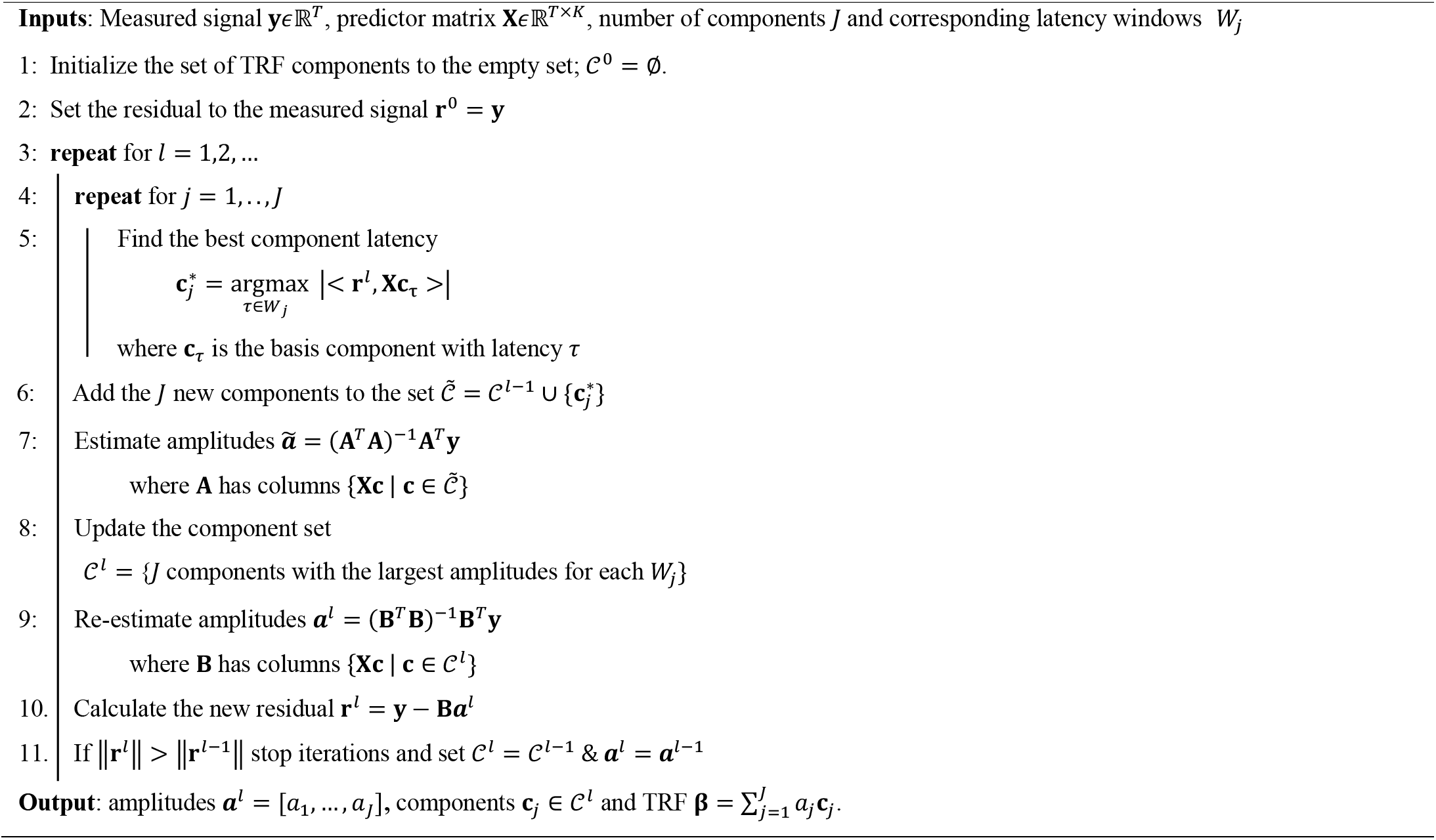

The SP algorithm estimates TRFs composed of only the required number of components, and can also be applied independently at each sensor for multi-channel TRFs.

### C. Proposed EM-SP algorithm for TRF Estimation

The EM-SP algorithm is an extension of the SP algorithm for multidimensional TRFs. In addition to directly estimating amplitudes and latencies, this algorithm also directly estimates sensor topographies or source distributions for multi-channel TRFs. This algorithm uses EM to iteratively estimate component topographies in the E-step, and latencies using SP in the M-step. Given a predefined number of components and corresponding latency windows, the EM-SP multi-channel TRF model is given by a modified version of (3).

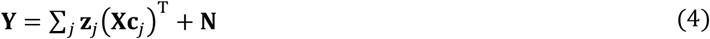

Where 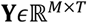 is the measured data over *M* sensors and *T* time points, 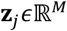 the spatial topography of the *j^th^* component, 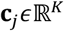 is the temporal waveform of the *j^th^* component, 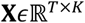 is the predictor matrix, and 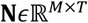 is the measurement noise. The component latency is given by *τ_j_* and is related to (4) by the fact that **c***_j_* corresponds to column number *τ_j_* in the TRF basis dictionary matrix. We assume the following priors,

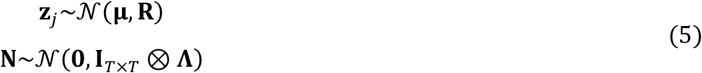

Where the temporal noise covariance is assumed to be the identity matrix and the spatial noise covariance is given by 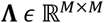. For the EM algorithm, we consider the spatial topographies 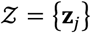 as the ‘hidden’ variables. The remaining parameters that need to be estimated are Θ = {**μ, R, Λ**, *τ_j_*}. Detailed derivations of the algorithm are provided in supplementary materials. Here, we summarize the main steps of the algorithm.

The Q-function is given by taking the expectation over the posterior probability 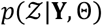.

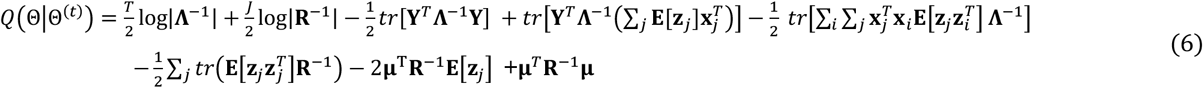

In the Expectation step, the posterior means of the spatial topographies are estimated.

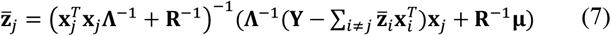

For the Maximization step, we use the Conditional Maximization method [32] whereby we sequentially maximize over each one of the parameters Θ = {**μ, R, Λ**, *τ_j_*,}, while holding the others fixed at their previous values.

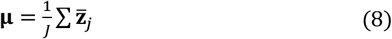

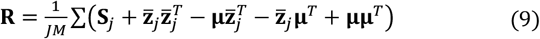

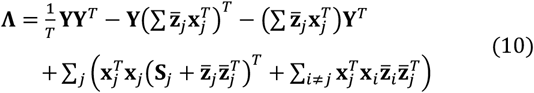

The latencies *τ_j_* can be estimated in a similar manner to the single channel SP algorithm using a linear search to maximize 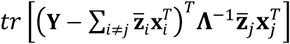 over the component basis. The complete EM-SP algorithm is provided below.

#### Algorithm 2: EM-SP

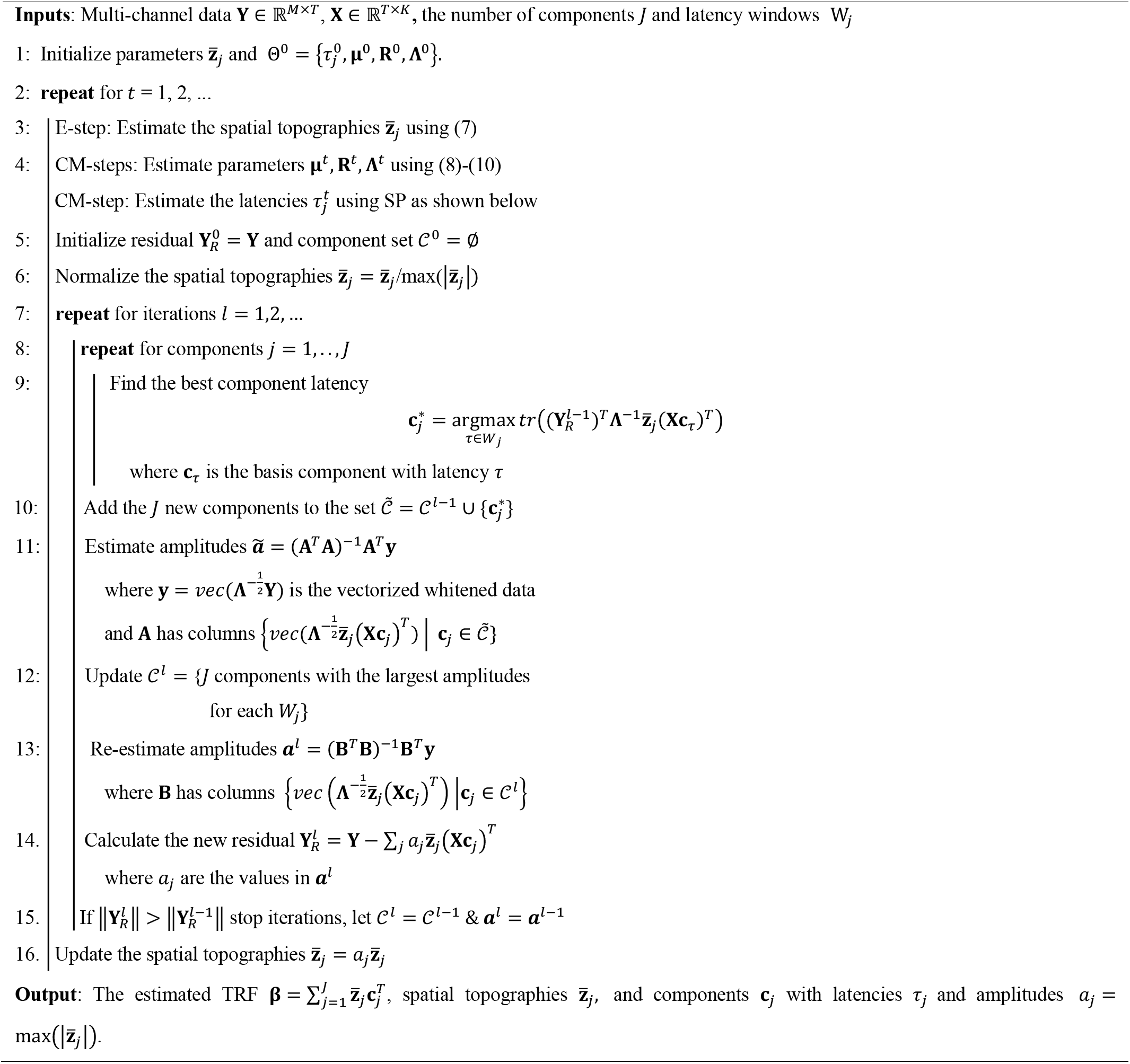

All four algorithms can also be used to simultaneously fit TRFs to multiple predictors (e.g., foreground and background envelopes) by concatenating the *P* predictor matrices 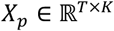 along the columns, resulting in a new predictor matrix 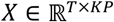. In this work, we jointly fit TRFs to two predictors (corresponding to foreground and background speech envelopes) using a concatenated predictor matrix.

### D. Simulation Study

Simulations were constructed to match typical cocktail party speech experiments which have two simultaneous speech streams. Accordingly, the envelopes of two speech stimuli (foreground and background) were used as predictors. These envelopes were constructed by first passing the speech waveform through a gammatone filterbank with 256 frequency bands between 20-4000 Hz, and the amplitude spectrogram was computed with an integration window of 10 ms. The resulting spectrogram was averaged across frequency bands, downsampled to 1000 Hz, and then band-passed at 1-10 Hz using a symmetric linear phase FIR filter with order 3301 and cutoffs 0.5 Hz and 11.25 Hz. Finally, the envelopes were downsampled to 100 Hz for all further analysis. These envelopes were repeated three times, in line with experiments having multiple trials of repeated stimuli to extract consistent responses using spatial filters such as Denoising Source Separation (DSS [33]; details given below). Each predictor was convolved with its own ground truth simulated TRF and the responses were summed together to form one-dimensional responses at 100 Hz sampling rate for 30 pseudo-subjects comparable to a single-sensor M/EEG response or the first auditory response component after DSS.

For each simulated subject, the ground truth simulated TRF was formed by placing Hamming windows of 50 ms width with peaks in the latency ranges 30-80 ms, 90-170 ms and 190-250 ms corresponding to typical latencies of the M50, M100 and M200 components. The M100 component was given a negative sign, and the components were scaled and shifted according to randomized subject specific amplitudes and latencies. These amplitudes and latencies were later used as the ground truth for performance evaluation.

Realistic noise was added to the simulated responses using the first DSS component of real MEG data collected from 30 subjects listening to speech (previously published [34], [35]). DSS creates a series of spatial filters, where the DSS component generated by the first of these filters corresponds to activity that is most consistent across repeated stimulus presentations (see [33] for details on DSS). Therefore, for this speech experiment, the first DSS component is dominated by auditory activity and displays a typical auditory response sensor topography. This component was then phase scrambled, preserving the spectral properties of MEG signals, to simulate noise added to the simulated response, at SNRs of −15, −20, −25 and −30 dB (SNRs selected to result in realistic TRF model correlation values).

The multi-channel simulation followed the same method for 157 simulated sensor signals, but in addition also used ground truth sensor topographies for each TRF component. These topographies were constructed from the TRF component topographies of a real subject with typical auditory TRF components, with the addition of Gaussian noise to simulate individual variability. Real multi-channel MEG data was again phase scrambled and added as noise on a per channel basis using the method described above, at SNRs of −20, −25, −30 and −35 dB (lower SNRs were used because unprocessed multi-channel data is typically noisier than the extracted auditory component).

The DSS algorithm was also applied to the simulated multi-channel data and corresponding TRFs were calculated for the first 6 DSS components. These DSS TRFs were projected back into sensor space for subsequent analysis and for computing performance metrics.

The source space simulation was constructed using the Freesurfer ico-4 surface source space of the ‘fsaverage’ brain [36]. An ROI in temporal lobe with 245 sources that included auditory cortex was used for this simulation (‘aparc’ labels ‘transversetemporal’ and ‘superiortemporal’). The three TRF components were simulated using dipoles in Heschl’s gyrus, Planum Temporale and Superior Temporal Gyrus in both hemispheres. These dipoles were projected onto the sensors using forward models from real data and back projected back onto source space with Minimum Norm Estimation (MNE) [37] using Eelbrain [14], [38] and MNE-Python softwares [39] to simulate the source localization procedure. The back-projected source distributions of these simulated TRF components were also used as the ground truth for subsequent performance comparisons. The TRFs were then convolved with the predictors to form the responses at each of the 245 sources. Real MEG data was phase scrambled and added as noise to the response at each source at SNRs of −15, −20, −25 and −30 dB following the same procedure as above.

### E. Experimental Dataset

MEG data collected in a prior study [34], [35] was used for evaluating the performance of the algorithms on real data. The study was approved by the IRB of the University of Maryland and all participants provided written informed consent prior to the start of the experiment. The dataset consisted of MEG data collected from 40 subjects while they listened to speech from the narration of an audiobook. Subjects listened to two speakers simultaneously in a cocktail party experiment, but were asked to attend to only one speaker. The data was from the condition where the foreground speaker was 3 dB louder than the background speaker. TRFs were estimated for the foreground and background envelopes. Whole head sensor space TRFs (157 sensors) were computed for each algorithm on three minutes of data. Additionally, TRFs were also computed for the first 6 DSS components. Finally, the MEG responses of this dataset were source localized using MNE and source space TRFs were also computed.

### F. Algorithm Implementation

The algorithms were implemented in Python (version 3.9.6) using SciPy (version 1.8.0) [40], and Eelbrain (version 0.36.1). The code is available online at <URLs available upon acceptance>. A basis dictionary with Hamming windows of width 50 ms was used for boosting, SP and EM-SP. The component latency windows for the SP and EM-SP algorithms were 30-80 ms, 90-170 ms and 190-250 ms. To avoid instability and convergence issues, the spatial covariance **R** for the EM-SP algorithm was assumed to be the identity matrix. The EM-SP was initialized using the extracted components from the SP algorithm applied at each sensor/source independently.

A nested 4-fold cross validation procedure was followed for all algorithms to allow for unbiased comparison. The data was divided into 4 splits, with 1 for testing, 1 for validation and 2 for training. The validation and training splits were permuted for each test split in a nested fashion. The training data was used to optimize the ridge TRF over several regularization parameters (steps of 2^0^, 2^1^, …, 2^16^) based on the model fit on the validation data. The boosting TRF was fit on the training data, and the validation data was used to check for convergence and terminate the algorithm. The SP and EM-SP TRFs were fit on the training data, and the model fit on the validation data was used to terminate the EM iterations. Finally, the overall model fit metric was calculated by convolving the average TRF over all training splits with the appropriate test predictor and computing the Pearson correlation between this predicted signal and the actual test signal.

### G. Performance Metrics

The model fit was calculated as the Pearson correlation between the estimated and the predicted response (averaged over channels for multidimensional cases). A null model was constructed by fitting TRFs using circularly time-shifted predictors (shifts of 15 s) and the correlation of this null model was subtracted from the true model. This bias corrected model fit is reported for both simulations and real data.

In addition to model fit, several other metrics of TRF component estimation were also calculated for the simulations (but not for real data, since the ground truth components were unknown). TRF components were automatically detected as the peaks of the r.m.s of the TRF across channels in the appropriate latency windows (30-80 ms, 90-170 ms, 190-250 ms) and the following metrics were used; 1) Pearson correlation between the estimated and ground truth TRF, 2) Absolute error of individual component latency estimates 3) Absolute error of individual component amplitude estimates (estimated vs, ground truth), 4) Spurious TRF activity given by the % r.m.s. power in the estimated TRF after 300 ms (note that there is no activity in the ground truth TRF after 300 ms), 5) Number of missing components 6) Sensor/source topography estimation error using the angle between the estimated topography vector and the ground truth topography vector. These metrics were averaged over channels, predictors, and components.

## III. Results

### A. Simulation: Single-Channel TRFs

Single-channel TRFs were simulated, and the ridge, boosting, and SP algorithms were compared in terms of several performance metrics. The estimated TRFs for a representative subject are shown in Fig. 1. The conventional measure for evaluating the performance of TRF models is the correlation between the actual and the predicted responses. In this work we used a nested cross-validation procedure for all algorithms to reduce overfitting and a null model based on shifted predictors for bias correction. However, correlation between the actual and the predicted responses may not always be an appropriate measure of TRF component estimation, since it depends on a variety of factors including SNR and predictor characteristics. This metric may also not appropriately penalize latency errors or spurious activity in the TRF. Hence, we used several other metrics, including component latency and amplitude errors, to compare these algorithms in terms of TRF component estimation (see right column of Fig. 1).

**Fig. 1.**
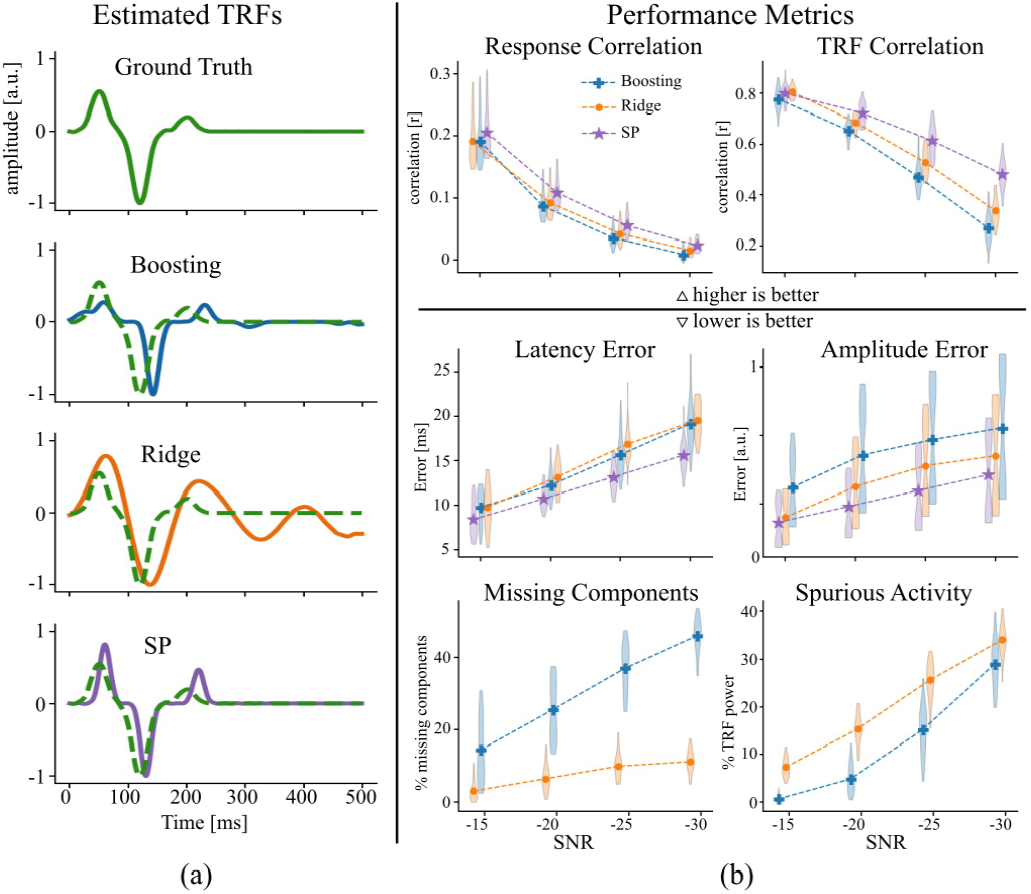
Performance comparison for single-channel simulations. (a) The fitted TRFs for a representative subject. The ground truth TRF is shown as a dotted green line over the estimated TRFs. (b) Algorithm comparison using the performance metrics. Violin plots over simulated subjects are shown, with the symbols indicating the mean. Within each SNR condition, the algorithms are plotted in ascending order of their means from left to right. SP does not have spurious activity after 300 ms or missing components by design and is not shown for the bottom two subplots. Boosting seems to miss some components, while ridge has more spurious activity. Ridge and boosting are comparable for most measures, while SP seems to outperform the others in higher SNR cases.

The SP algorithm performed the best in most measures, while ridge and boosting performed comparably. Spurious peaks after 300 ms (when there was no activity in the ground truth TRF) could lead to difficulties in interpretation and to false positives when detecting TRF components in real data. Conversely, missing components (false negatives) could also lead to improper interpretation of TRFs. Ridge had more spurious activity than boosting but was also able to detect more components than boosting.

### B. Simulation: Multi-channel TRFs

Sensor space TRFs were simulated using realistic sensor topographies for TRF components, and the performance of each algorithm was compared (see Fig. 2). TRFs were estimated independently at each sensor for the boosting, ridge and SP algorithms, while the EM-SP algorithm directly estimated multi-channel component topographies. The EM-SP algorithm performed the best in most measures, while ridge and boosting performed comparably. The sensor topographies estimated by boosting and SP are worse than those estimated by ridge and EM-SP, which is to be expected given that the former are sparse algorithms that are fit at each sensor independently. Interestingly, the missing components are similar for both ridge and boosting, unlike in the single-channel case. If boosting is able to correctly estimate components even for only a few channels, sparsity (in time) can then preserve the presence of the component peak when the r.m.s of the TRF is taken across channels. This improvement in component detection for boosting is also seen for the DSS and source space TRFs reported below.

**Fig. 2.**
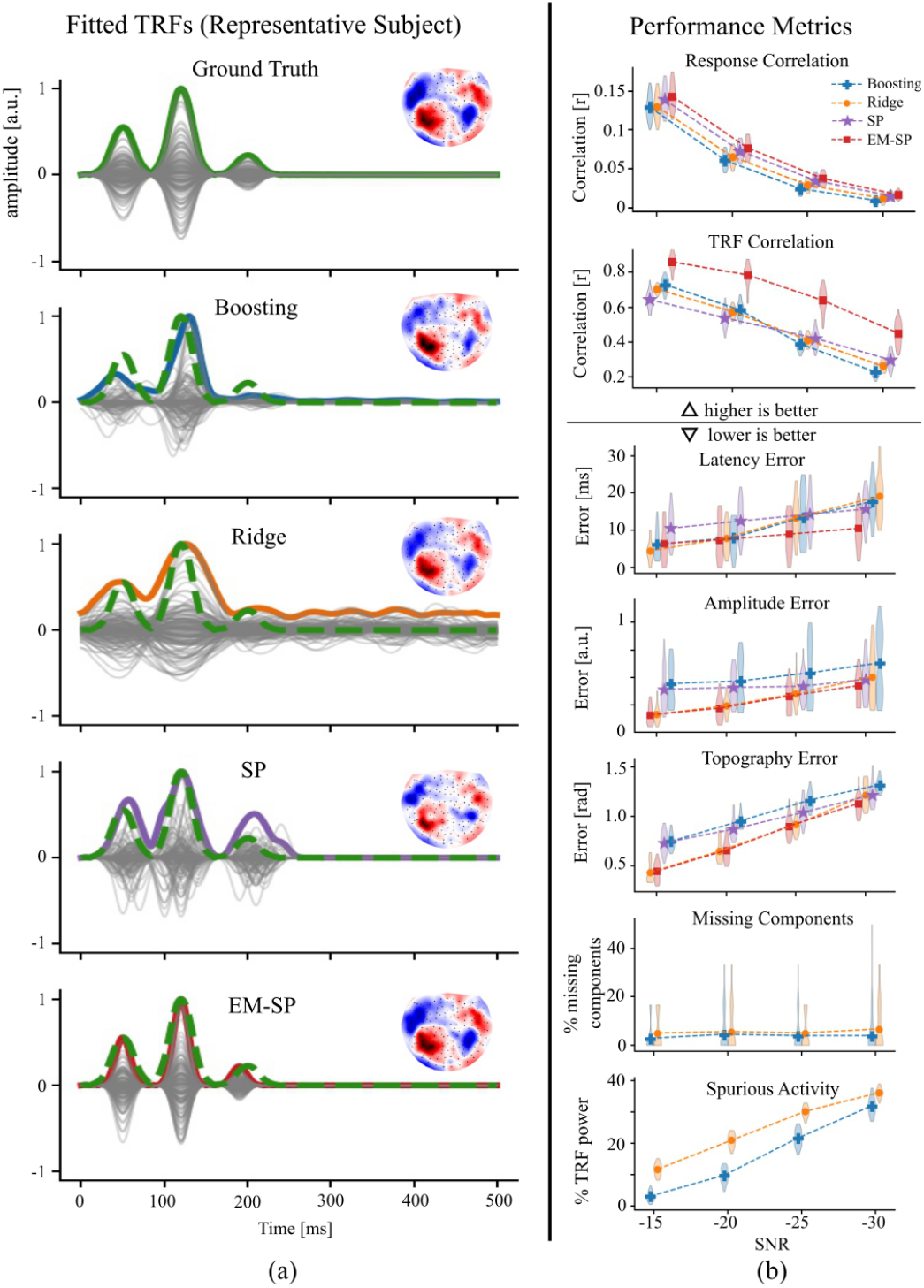
Performance comparison for multi-channel simulations. (a) The fitted TRFs for a representative subject. The TRF at each sensor is plotted in gray, while the ℓ_2_-norm over sensors is plotted as a colored thick line. The ℓ_2_-norm of the ground truth TRF is shown as a dotted green line over the estimated TRFs. The sensor topography at the largest peak near 100 ms is shown as an inset. (b) Algorithm comparison using the performance metrics. Since there is no activity after 300 ms in the SP and EM-SP TRFs by design, they are not plotted in the spurious activity subplot. EM-SP outperforms the others in most measures. Although all methods find similar components, the sensor topographies for boosting and SP are worse than the others, perhaps because they are sparse estimation techniques.

### C. Simulation: Denoised TRFs using DSS

The DSS algorithm was applied to the simulated sensor space responses to extract spatial filters corresponding to auditory response components. The algorithms were fit on the first 6 DSS components, and the resulting TRFs were projected back onto the sensor space for performance evaluation. Model fit response correlations increased greatly over the sensor space TRFs in all cases (see Fig. 3). Ridge, boosting and EM-SP had comparable results. Interestingly, EM-SP did not have a significant advantage over the other algorithms, indicating that the established algorithms are just as suitable for low dimensional, denoised data.

**Fig. 3.**
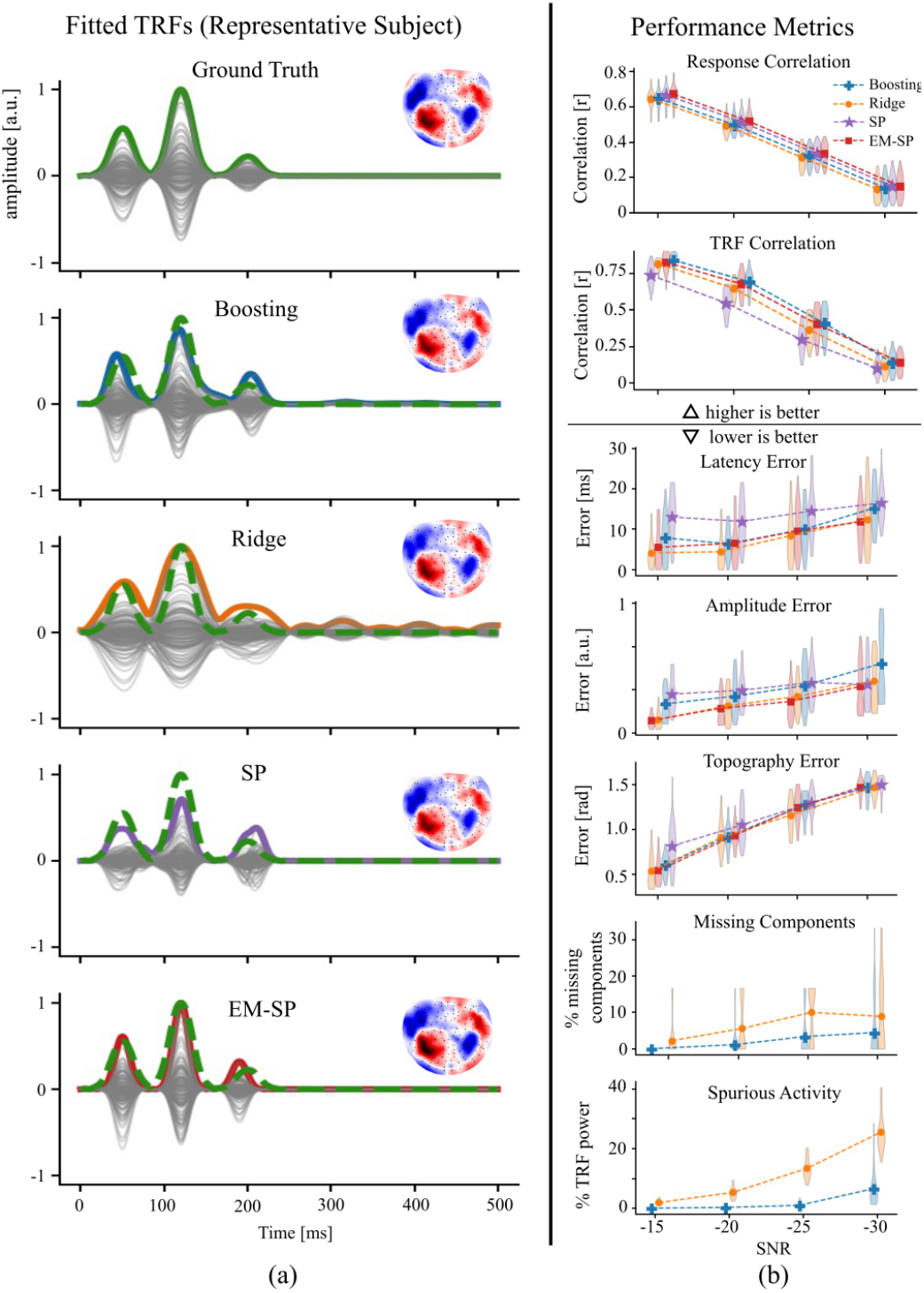
Performance comparison after DSS denoising. (a). The fitted TRFs for a representative subject, similar to the previous figure. The TRFs were fit on the first 6 DSS components and then back-projected to sensor space. All the algorithms result in reasonable TRF components and sensor topographies. (b). Algorithm comparison using the performance metrics. All the algorithms except SP perform comparably, while the latter performs the worst in most cases.

### D. Simulation: Source Localized TRFs

Source space simulations were constructed with dipoles in auditory areas for each TRF component. These dipoles were projected onto sensor space using the forward model and source localized back to source space to simulate source localized MEG data. The algorithms were fit on these source localized signals and performance was compared using the same metrics (see Fig. 4). Results were similar to the sensor space simulation, with EM-SP outperforming the others, and ridge and boosting giving comparable results (with ridge typically performing marginally better than boosting for most measures except spurious activity).

**Fig. 4.**
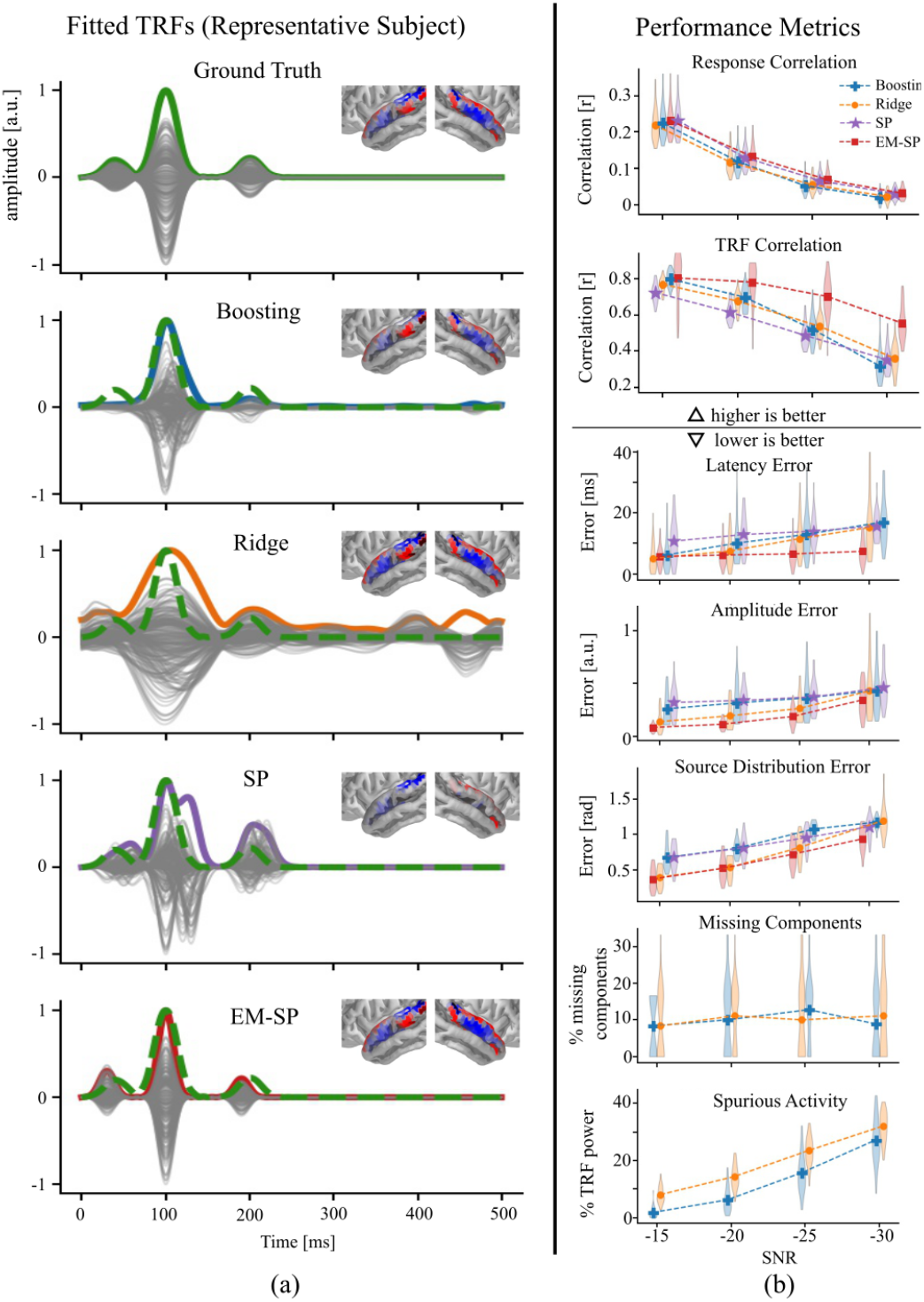
Performance comparison for source space simulations. (a) The fitted TRFs for a representative subject are shown, similar to the previous figure. The source distributions in the temporal lobe ROI at the largest peak near 100 ms are shown as insets. Boosting and SP result in much sparser source distributions, and all the algorithms except SP perform comparably in estimating the TRF components, although the ridge TRF has a lot more activity that may make it difficult to interpret in realistic situations where the ground truth is unknown. (b). Algorithm comparison using the performance metrics, similar to those shown in the previous figure. EM-SP outperforms the others in most cases.

Overall, the simulation results indicate that both boosting and ridge are comparable, with ridge typically performing slightly better. Interestingly, SP outperformed ridge and boosting in the high noise single-channel simulations, while EM-SP outperformed the others by a large margin in the multi-channel and source-localized simulations. It should be noted that the component windows used for the simulation were identical to the component windows provided a-priori to SP and EM-SP, which may explain their better performance. Therefore, SP and EM-SP may be suitable for estimating TRFs in high noise conditions, assuming that the appropriate latency windows can be determined a-priori. Ridge also had lower spatial error compared to boosting (sensor topography and source distribution errors), perhaps because a sparse estimation technique like boosting cannot capture smooth spatial patterns as well as ridge. Conversely, ridge had much larger amounts of spurious activity compared to boosting. However, after applying the DSS algorithm, ridge, boosting and EM-SP once again showed comparable performance, highlighting the importance of denoising methods when estimating TRFs from noisy multidimensional data.

### E. Performance on Real Data

The algorithms were compared on a real MEG dataset collected for a cocktail party experiment. Sensor space, DSS and source space TRFs are shown for a representative subject in Fig. 5. The only metric used was the correlation between the measured and predicted signals, since the other metrics cannot be calculated when the ground truth TRF components are unknown. Interestingly, boosting had significantly lower correlation accuracy compared to each of the three other algorithms for sensor and source space TRFs (paired samples permutation tests with Holm-Bonferroni correction; all comparisons with boosting resulted in t39 > 4, p < 0.01), but there were no significant differences in correlation accuracy between ridge, SP and EM-SP. However, it is unclear if correlation is the most suitable metric for evaluating the accuracy of estimating TRF components. The correlation values were distributed over a large range across subjects, possibly indicating a high degree of inter-subject variability in neural SNR for time-locked responses. Ridge resulted in smooth TRFs with several peaks and large amounts of non-zero activity which made them more difficult to interpret, especially for the sensor and source space TRFs. Boosting, though performing worse in terms of correlation, allowed for sparser TRFs with fewer peaks that were easier to interpret.

**Fig. 5.**
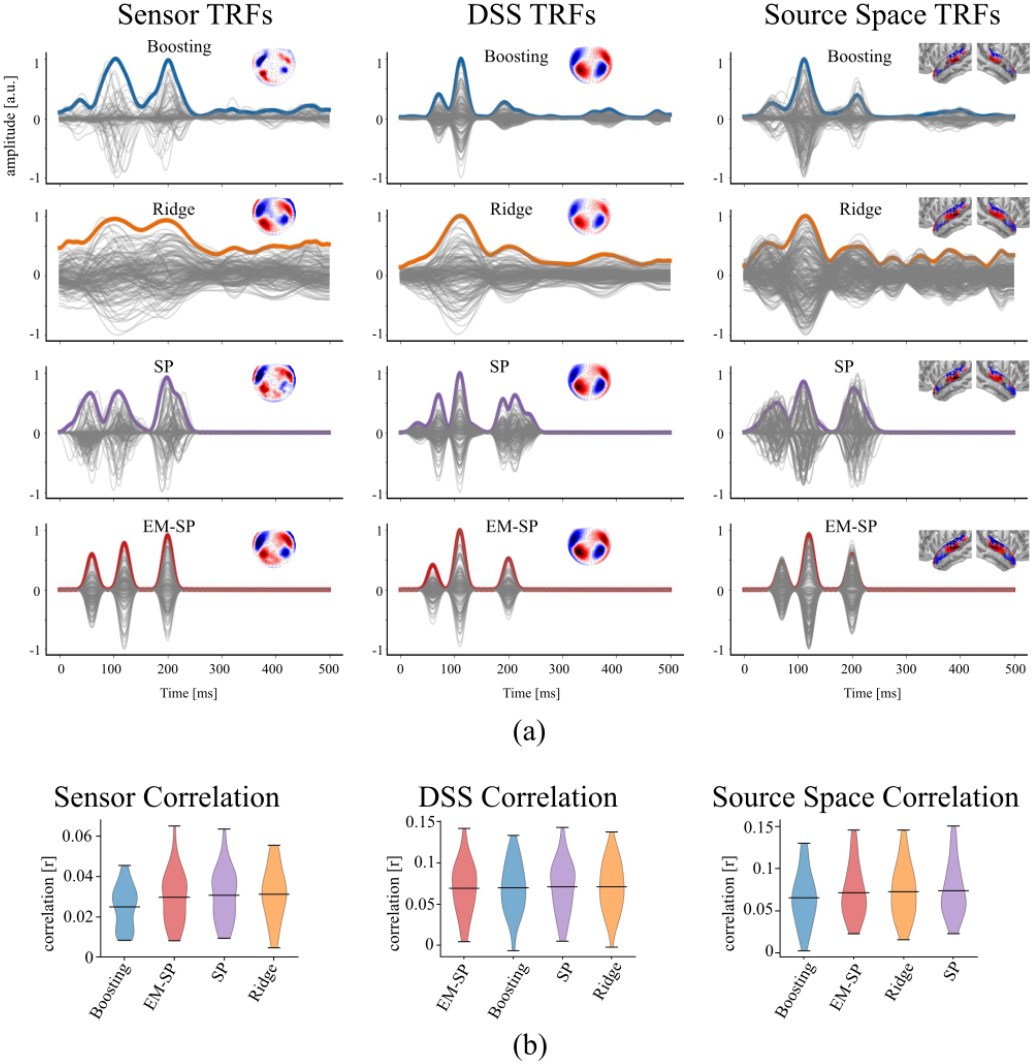
Performance comparison on real MEG data. (a) The estimated sensor, DSS and source localized TRFs are shown for a representative subject. The sensor topographies and source distributions at the large peak near 100 ms are shown as insets. The sensor space EM-SP TRF has clear components and topographies, while the boosting TRF has overly sparse topographies and the ridge TRF has a lot of hard to interpret activity. Boosting, ridge and EM-SP show clear components and spatial patterns for the DSS and source localized TRFs. (b) Correlation between the measured and predicted signals is shown as a measure of model fit. Violin plots across subjects are shown for each algorithm in ascending order of their mean from left to right.

The two proposed algorithms were restricted to finding exactly three TRF components, assuming fixed component waveforms and latency windows. The fact that EM-SP may have performed worse than ridge for real data, even though it outperformed the others in the simulations, indicates that these assumptions may not be valid for all subjects. This could be due to a variety of reasons including missing components due to anatomical or functional differences, and large individual variability in TRF components latencies, waveforms, and peak widths. Indeed, a separate simulation analysis (not shown) with missing components and mismatched latency windows resulted in similar performance for EM-SP, with it no longer outperforming ridge and boosting. In any case, conventional post-hoc analysis of TRF components estimated using established algorithms is also typically performed under similar assumptions to those used for EM-SP (i.e., detecting TRF peaks using similar latency windows). However, even with these constraints, EM-SP was often able to recover TRF components and spatial patterns comparable to ridge.

## IV. Conclusion

TRFs provide a significant advancement over ERPs, allowing for experiments with more naturalistic speech paradigms. Detecting robust TRF components is essential for reliable single-subject investigations that could inform diagnosis and treatment of hearing disabilities and lead to improved biomedical applications like smart hearing aids.

We compared TRF algorithms using both model fit and component estimation accuracy. Simulations indicate that boosting and ridge are comparable for most cases. Interestingly, ridge had better model fits on real data. However, in general, ridge TRFs displayed more spurious activity, while boosting TRF peaks were more interpretable. Therefore, ridge may be suitable for studies focused on prediction accuracy, while boosting may be appropriate for detecting easily identifiable TRF components. We restricted our analysis of established methods to these two algorithms that are the most widely used. Other variations on regularized regression, such as Lasso and Elastic Net, may provide improvements in TRF estimation [12].

SP and EM-SP performed exceptionally in simulations, but not on real data, possibly due to invalid assumptions. The a-priori parameters may need to be tuned for each predictor type or experiment, or even for each subject

Modern TRF analyses involve multiple types of predictors [42] (e.g., envelopes, phoneme onsets, multiple frequency bands for spectrotemporal TRFs). Boosting and banded ridge regression may be suitable for these studies [10], [13], [43], [44]. The component characteristics of TRFs to these higher-level predictors must be determined before SP and EM-SP can be applied. Additionally, early low-level responses could impact TRFs to high-level predictors, and sparse algorithms with fewer false positives (but more false negatives) may be more conservative.

In conclusion, our results indicate that SP and EM-SP may only perform well under realistic assumptions, while ridge and boosting perform comparably in most cases, with ridge typically having higher prediction accuracies, but also more spurious activity.

