## Supplementary Materials for "Algorithms for Estimating Time-Locked Neural Response Components in Cortical Processing of Continuous Speech"

### The EM-SP Algorithm for TRF Estimation

Mathematical derivations for the EM-SP algorithm are provided here. This algorithm uses the Expectation Maximization (EM) method to iteratively estimate TRF component amplitudes and topographies in the E-step, and latencies using SP in the M-step. Given a predefined number of components and corresponding latency windows, the multichannel TRF model is given by.

$$\mathbf{Y} = \sum_j \mathbf{z}_j \left(\mathbf{X}\mathbf{c}_j\right)^{\mathrm{T}} + \mathbf{N} \tag{S1}$$

Where $\mathbf{Y}\epsilon\mathbb{R}^{M\times T}$ is the measured data over $M$ sensors and $T$ time points, $\mathbf{z}_j\epsilon\mathbb{R}^{M}$ is the spatial topography of the $j^{th}$ component, $\mathbf{c}_j\epsilon\mathbb{R}^{K}$ is the temporal waveform of the $j^{th}$ component and $\mathbf{N}\epsilon\mathbb{R}^{M\times T}$ is the measurement noise. $\mathbf{X}\epsilon\mathbb{R}^{T\times K}$ is the predictor matrix with each column corresponding to lagged predictors. The component latency is given by $\tau_j$ and is related to (S1) by the fact that $\boldsymbol{c}_j$ corresponds to column number $\tau_j$ in the TRF basis dictionary matrix. We assume the following priors,

$$\begin{gathered}\mathbf{z}_j \sim \mathcal{N}(\boldsymbol{\mu}, \mathbf{R}) \\ \mathbf{N} \sim \mathcal{N}(\mathbf{0}, \mathbf{I}_{T\times T} \otimes \boldsymbol{\Lambda})\end{gathered} \tag{S2}$$

Where the temporal noise covariance is assumed to be the identity matrix and the spatial noise covariance is given by $\boldsymbol{\Lambda}\epsilon\mathbb{R}^{M\times M}$. For the EM algorithm, we consider the spatial topographies $\mathcal{Z} = \{\mathbf{z}_j\}$ as the 'hidden' variables. The remaining parameters that need to be estimated are $\Theta = \{\tau_j, \boldsymbol{\mu}, \mathbf{R}, \boldsymbol{\Lambda}\}$. The data likelihood is given by

$$p(\mathbf{Y}|\mathcal{Z};\Theta) \sim \mathcal{N}\left(\sum_j \mathbf{z}_j \mathbf{x}_j^T, \mathbf{I}_{T\times T} \otimes \boldsymbol{\Lambda}\right) \tag{S3}$$

Where for convenience we have denoted $\mathbf{x}_j^T = \left(\mathbf{X}\mathbf{c}_j\right)^T$. Assuming $\{\mathbf{z}_j\}$ are i.i.d., the complete data log likelihood is then given by

$$\begin{aligned}\mathcal{L}_c(\mathbf{Y},\mathcal{Z};\Theta) &= \log p(\mathbf{Y}|\mathcal{Z};\Theta) + \sum_j \log p(\mathbf{z}_j) \\ &\propto \frac{T}{2}\log|\boldsymbol{\Lambda}^{-1}| - \frac{1}{2} tr\left[\left(\mathbf{Y} - \sum_j \mathbf{z}_j\mathbf{x}_j^T\right)^T \boldsymbol{\Lambda}^{-1}\left(\mathbf{Y} - \sum_j \mathbf{z}_j\mathbf{x}_j^T\right)\right] \\ &\quad + \frac{J}{2}\log|\mathbf{R}^{-1}| - \frac{1}{2}\sum_j\left(\mathbf{z}_j - \boldsymbol{\mu}\right)^T \mathbf{R}^{-1}\left(\mathbf{z}_j - \boldsymbol{\mu}\right)\end{aligned} \tag{S4}$$

Hence, the Q-function is given by

$$\begin{aligned}Q\left(\Theta|\Theta^{(k)}\right) &= \mathbf{E}_{\mathcal{Z}|\mathbf{Y};\Theta}[\mathcal{L}_c(\mathbf{Y},\mathcal{Z};\Theta)] \\ &= \frac{T}{2}\log|\boldsymbol{\Lambda}^{-1}| - \frac{1}{2} tr[\mathbf{Y}^T\boldsymbol{\Lambda}^{-1}\mathbf{Y}] + tr\left[\mathbf{Y}^T\boldsymbol{\Lambda}^{-1}\left(\sum_j \mathbf{E}[\mathbf{z}_j]\mathbf{x}_j^T\right)\right] \\ &\quad - \frac{1}{2} tr\left[\left(\sum_j\sum_i \mathbf{x}_j^T\mathbf{x}_i\mathbf{E}[\mathbf{z}_j\mathbf{z}_i^T]\right)\boldsymbol{\Lambda}^{-1}\right] \\ &\quad + \frac{J}{2}\log|\mathbf{R}^{-1}| - \frac{1}{2}\sum_j\left(tr(\mathbf{E}[\mathbf{z}_j\mathbf{z}_j^T]\mathbf{R}^{-1}) - 2\boldsymbol{\mu}^{\mathrm{T}}\mathbf{R}^{-1}\mathbf{E}[\mathbf{z}_j] + \boldsymbol{\mu}^T\mathbf{R}^{-1}\boldsymbol{\mu}\right)\end{aligned} \tag{S5}$$

The expectation is over the posterior distribution $p(\mathcal{Z}|\,\widetilde{\mathbf{Y}};\Theta) \propto \mathcal{L}_c(\mathbf{Y},\mathcal{Z};\Theta)$. Since this is quadratic in $\mathbf{z}_j$, the posterior for each $\mathbf{z}_j$ is normal.

$$p(\mathbf{z}_j|\,\widetilde{\mathbf{Y}};\Theta) = \mathcal{N}(\bar{\mathbf{z}}_j, \mathbf{S}_j) \tag{S6}$$

Using the properties of a Gaussian pdf, the mean $\bar{\mathbf{z}}_j$ of the Gaussian is given by setting the derivative to zero.

$$\mathbf{\Lambda}^{-1}(\mathbf{Y} - \textstyle\sum_i \mathbf{z}_i \mathbf{x}_i^T)\mathbf{x}_j - \mathbf{R}^{-1}(\mathbf{z}_j - \boldsymbol{\mu}) = 0 \tag{S7}$$

The covariance $\mathbf{S}_j$ of the Gaussian is given by the inverse of the Hessian.

$$\mathbf{S}_j = \left(\mathbf{x}_j^T \mathbf{x}_j \mathbf{\Lambda}^{-1} + \mathbf{R}^{-1}\right)^{-1} \tag{S8}$$

Using Eq. 5.12 and Eq. 5.13 the mean is given by

$$\bar{\mathbf{z}}_j = \mathbf{S}_j(\mathbf{\Lambda}^{-1}\widetilde{\mathbf{Y}}\mathbf{x}_j + \mathbf{R}^{-1}\boldsymbol{\mu}) \tag{S9}$$

$$where \quad \widetilde{\mathbf{Y}} = \mathbf{Y} - \textstyle\sum_{i \neq j} \mathbf{z}_i \mathbf{x}_i^T \tag{S10}$$

Note that the solution for each $\bar{\mathbf{z}}_j$ are coupled through $\widetilde{\mathbf{Y}}$ and a system of linear equations must be solved. However, in practice, solving each $\bar{\mathbf{z}}_j$ while keeping the others fixed leads to an approximate solution and avoids instabilities and computations of large matrix inversions. Therefore, we use (S9) for each $\bar{\mathbf{z}}_j$ to solve for the posterior mean. The relevant terms in the Q-function can now be substituted as follows

$$\mathbf{E}[\mathbf{z}_j] = \bar{\mathbf{z}}_j \tag{S11}$$

$$\mathbf{E}[\mathbf{z}_j \mathbf{z}_j^T] = \mathbf{S}_j + \bar{\mathbf{z}}_j \bar{\mathbf{z}}_j^T \tag{S12}$$

$$\mathbf{E}[\mathbf{z}_j \mathbf{z}_i^T] = \bar{\mathbf{z}}_j \bar{\mathbf{z}}_i^T \quad for\ i \neq j \tag{S13}$$

For the M-step, we maximize the Q-function w.r.t to the other parameters $\Theta = \{\tau_j, \boldsymbol{\mu}, \mathbf{R}, \mathbf{\Lambda}\}$. We use the Conditional Maximization method (Meng and Rubin, 1993) whereby we sequentially maximize over each one of these parameters while holding the others fixed at their previous values. Maximization updates are found by setting the partial derivatives of the Q-function to zero. For $\boldsymbol{\mu}$, the relevant terms of the Q-function are:

$$\begin{aligned} &-\tfrac{1}{2}\textstyle\sum_j\left(-2\boldsymbol{\mu}^{\mathrm{T}}\mathbf{R}^{-1}\mathbf{E}[\mathbf{z}_j] + \boldsymbol{\mu}^T\mathbf{R}^{-1}\boldsymbol{\mu}\right) \\ &= -\tfrac{1}{2}\textstyle\sum_j\left(-2\boldsymbol{\mu}^{\mathrm{T}}\mathbf{R}^{-1}\bar{\mathbf{z}}_j + \boldsymbol{\mu}^T\mathbf{R}^{-1}\boldsymbol{\mu}\right) \end{aligned} \tag{S14}$$

Setting the derivative to zero, we find the update

$$\boldsymbol{\mu} = \tfrac{1}{J}\textstyle\sum_j \bar{\mathbf{z}}_j \tag{S15}$$

For **R**, the relevant terms of the Q-function are:

$$\frac{J}{2}\log|\mathbf{R}^{-1}| - \frac{1}{2}\sum_j\left(tr\left((\mathbf{S}_j + \bar{\mathbf{z}}_j\bar{\mathbf{z}}_j^T)\mathbf{R}^{-1}\right) - \boldsymbol{\mu}^T\mathbf{R}^{-1}\bar{\mathbf{z}}_j - \bar{\mathbf{z}}_j^T\mathbf{R}^{-1}\boldsymbol{\mu} + \boldsymbol{\mu}^T\mathbf{R}^{-1}\boldsymbol{\mu}\right) \tag{S16}$$

Hence, setting the derivative w.r.t. $\mathbf{R}^{-1}$ to zero,

$$\mathbf{R} = \frac{1}{JM}\sum_j\left(\mathbf{S}_j + \bar{\mathbf{z}}_j\bar{\mathbf{z}}_j^T - \boldsymbol{\mu}\bar{\mathbf{z}}_j^T - \bar{\mathbf{z}}_j\boldsymbol{\mu}^T + \boldsymbol{\mu}\boldsymbol{\mu}^T\right) \tag{S17}$$

For **Λ,** the relevant terms of the Q-function are:

$$\begin{aligned}&\frac{T}{2}\log|\boldsymbol{\Lambda}^{-1}| - \frac{1}{2}tr[\mathbf{Y}^T\boldsymbol{\Lambda}^{-1}\mathbf{Y}] + \frac{1}{2}tr\left[\mathbf{Y}^T\boldsymbol{\Lambda}^{-1}\left(\sum_j\bar{\mathbf{z}}_j\mathbf{x}_j^T\right)\right] + \frac{1}{2}tr\left[\left(\sum_j\bar{\mathbf{z}}_j\mathbf{x}_j^T\right)^T\boldsymbol{\Lambda}^{-1}\mathbf{Y}\right]\\ &-\frac{1}{2}\sum_j\left(\mathbf{x}_j^T\mathbf{x}_j tr\left[(\mathbf{S}_j + \bar{\mathbf{z}}_j\bar{\mathbf{z}}_j^T)\boldsymbol{\Lambda}^{-1}\right] + \sum_{i\neq j}\mathbf{x}_j^T\mathbf{x}_i tr\left[\bar{\mathbf{z}}_j\bar{\mathbf{z}}_i^T\boldsymbol{\Lambda}^{-1}\right]\right)\end{aligned} \tag{S18}$$

Calculating the derivative w.r.t. $\boldsymbol{\Lambda}^{-1}$ using the same methods and equating to zero:

$$\boldsymbol{\Lambda} = \frac{1}{T}\left[\mathbf{Y}\mathbf{Y}^T - \mathbf{Y}\left(\sum_j\bar{\mathbf{z}}_j\mathbf{x}_j^T\right)^T - \left(\sum_j\bar{\mathbf{z}}_j\mathbf{x}_j^T\right)\mathbf{Y}^T + \sum_j\left(\mathbf{x}_j^T\mathbf{x}_j\left(\mathbf{S}_j + \bar{\mathbf{z}}_j\bar{\mathbf{z}}_j^T\right)^T + \sum_{i\neq j}\mathbf{x}_j^T\mathbf{x}_i\bar{\mathbf{z}}_i\bar{\mathbf{z}}_j^T\right)\right] \tag{S19}$$

For $\tau_j$**,** the relevant terms involving $\mathbf{x}_j$ in the Q-function are

$$tr\left[\widetilde{\mathbf{Y}}^T\boldsymbol{\Lambda}^{-1}\bar{\mathbf{z}}_j\mathbf{x}_j^T\right] - \frac{1}{2}\left[\mathbf{x}_j^T\mathbf{x}_j tr\left[\left(\mathbf{S}_j + \bar{\mathbf{z}}_j\bar{\mathbf{z}}_j^T\right)\boldsymbol{\Lambda}^{-1}\right] + \sum_{i\neq j}\mathbf{x}_j^T\mathbf{x}_i tr\left[\bar{\mathbf{z}}_j\bar{\mathbf{z}}_i^T\boldsymbol{\Lambda}^{-1}\right]\right] \tag{S20}$$

We assume that the second term doesn't depend on $\tau_j$ (i.e., $\mathbf{x}_j^T\mathbf{x}_j$ and $\mathbf{x}_j^T\mathbf{x}_i$ are similar for all rows of the lagged predictor matrix since they have similar vector norms). Therefore, the only term that depends on the latency is the first term $tr\left[\widetilde{\mathbf{Y}}^T\boldsymbol{\Lambda}^{-1}\bar{\mathbf{z}}_j\mathbf{x}_j^T\right]$, which is the correlation between the whitened measurements and predictions. To maximize this term, we use the SP algorithm, with appropriate modifications to include spatial topographies. The complete algorithm is provided in the paper.
